# Screening and functional analysis of differentially expressed genes in an animal model of *EBV*-associated lymphomas

**DOI:** 10.1101/666917

**Authors:** Yang Zhang, Chengkun Wang, Liangzhuan Liu, Qiu Peng, Xiaoning Gan, Lu Xie, Meiying Tan, Rongfang He, Yunlian Tang, Yimou Wu, Runliang Gan

## Abstract

*Epstein-Barr virus* (EBV) is an important human oncogenic virus. This paper is to explore how EBV induce malignant transformation of human lymphocytes and the related mechanism of lymphomagenesis. We have constructed *hu-PBL/SCID chimeric mice* and established a model of EBV-associated human-derived lymphomas. By using Agilent human whole genome microarray and a series of bioinformatic analyses, a total of 202 differentially expressed genes were screened from the EBV-induced lymphomas in *hu-PBL/SCID mice*, including 44 up-regulated and 158 down-regulated genes. Calculation of the rank score (RS) values of these genes in the HIPPIE protein interaction networks showed that topoisomerase II alpha (TOP2A), ubiquitin like with PHD and ring finger domains 1 (UHRF1), histone cluster 2 H2B family member E (HIST2H2BE), phosphoglycerate dehydrogenase (PHGDH), vinculin (VCL), insulin-like growth factor 1 receptor (IGF1R), Fos proto-oncogene (FOS), snail family transcriptional repressor 1 (SNAI1), PDZ binding kinase (PBK), and ring finger protein 144B (RNF144B) were the top 10 key node genes of EBV-induced lymphoma. In which, PBK, an up-regulated genes with the highest number of GO annotations, was verified by cellular function experiments and clinical lymphoma samples.

**Author summary:** EB virus is closely associated with human lymphoma and nasopharyngeal carcinoma. Since the susceptible hosts of EBV limit to human and cottontop tammarins, there are no appropriate animal models so far to study the EBV-associated oncogenesis. In our previous experiments, the EBV-associated lymphomas were induced in *hu-PBL/SCID chimera* (a new humanized mouse model). However, the cellular and molecular mechanisms of malignant transformation of normal human cells and tumor formation induced by EBV remain unclear. In this study, we examined and compared the gene expression profiles of EBV-induced lymphomas and normal human lymphocytes of the same origin in SCID mice. By constructing the gene-function relationship network, we preliminarily found that TOP2A, UHRF1, HIST2H2BE, PHGDH, VCL, IGF1R, FOS, SNAI1, PBK, and RNF144B may be the key genes in EBV-induced lymphomas. These findings suggest that the induction of lymphoma by EBV is a complex process that involves multiple genes and pathways.

## Introduction

*Epstein-Barr virus (EBV)* is an important human oncogenic virus that is closely associated with several human malignancies, including nasopharyngeal carcinoma and lymphoma. Analyses of clinic-pathological specimens have confirmed *EBV* infection in 55%∼80% cases of Hodgkin lymphomas [1,2] and 10∼42% cases of non-Hodgkin lymphomas [3,4]. Furthermore, *EBV* infection is detected in 90%∼100% of nasal/nasopharyngeal natural killer/T-cell lymphomas [5,6], while latent *EBV* infection is found in almost all Burkitt lymphoma among Africans [7]. It has been reported that *EBV*-positive lymphoma patients have a poorer prognosis than *EBV*-negative lymphoma patients [8,9].

*In vitro* experiments have demonstrated that *EBV* can transform and even immortalize normal human peripheral lymphocytes [10]. Our group was among the first to transplant *EBV*-infected human peripheral lymphocytes (hu-PBL) from healthy blood donors into *severe combined immunodeficiency (SCID) mice* to induce the generation of human-derived B cell lymphomas [11,12]. However, the cellular and molecular mechanisms of malignant transformation of normal lymphocytes and tumor formation induced by *EB virus* remain unclear.

Oncogenesis of virus-infected host cells involves not only specific gene products or protein molecules, but also the multidimensional functional interaction network between numerous genes and their expression products, which regulates the biological behaviors and characteristics of cells. There is no doubt that the expression and function of many genes are altered during *EBV* infection and its interactions with the cells; therefore, strategy of the single gene analysis is no longer adequate to investigate the complex molecular regulatory mechanism involved in this process. In our present study, the gene expression profiles of *EBV*-induced lymphomas in *hu-PBL/SCID chimeric mice* and normal human lymphocytes of the same origin were examined and compared using a human genome microarray to analyze differential gene expression in host cells before and after the development of *EBV*-induced lymphoma by bioinformatics. A molecular network and key nodes for *EBV*-induced lymphomas were constructed to screen for key genes and to provide new insights into the oncogenic mechanism of *EB virus*.

## Results

### Construction of *hu-PBL/SCID chimeric mice* and identification of *EBV*-induced tumors

*EBV*-associated tumors were developed in the *SCID mice* grafted with PBLs isolated from the six blood donors (Supplementary Table 1). Tumors were visible in the abdominal and/or thoracic cavity of the nine *SCID mice*, but since these mice received PBLs from six blood donors, the number of *EBV*-induced tumors was considered to be six as well. Macroscopic examination revealed irregular and nodular solid tumors with the larger ones at 1.6 × 1.5 × 1.0 cm^3^ and smaller ones at 0.7 × 0.5 × 0.5 cm^3^.

Microscopic observation of the induced tumors showed that the tumor cells were relatively large and were a mixture of cells with large fissure and without fissure. These cells have large cytoplasm that stained light red with plasma cell-, centroblast- and immunoblast-like morphologies (Fig 1A). Immunohistochemical staining indicated that the induced tumor cells were LCA-positive, CD20/CD79a (B cell marker)-positive, and CD3/CD45RO (T cell marker)-negative. These pathological characteristics of the induced tumor cells were consistent with those of diffuse large B-cell lymphoma. PCR amplification of the human-specific Alu sequence from the induced tumor tissues showed that the 221-bp amplicon of the Alu sequence was present in all six induced tumors, which was consistent with the positive control (Fig 1B), indicating that the induced tumors in the hu-PBL/SCID chimeric mice were of human origin and not mouse origin.

**Fig1.**
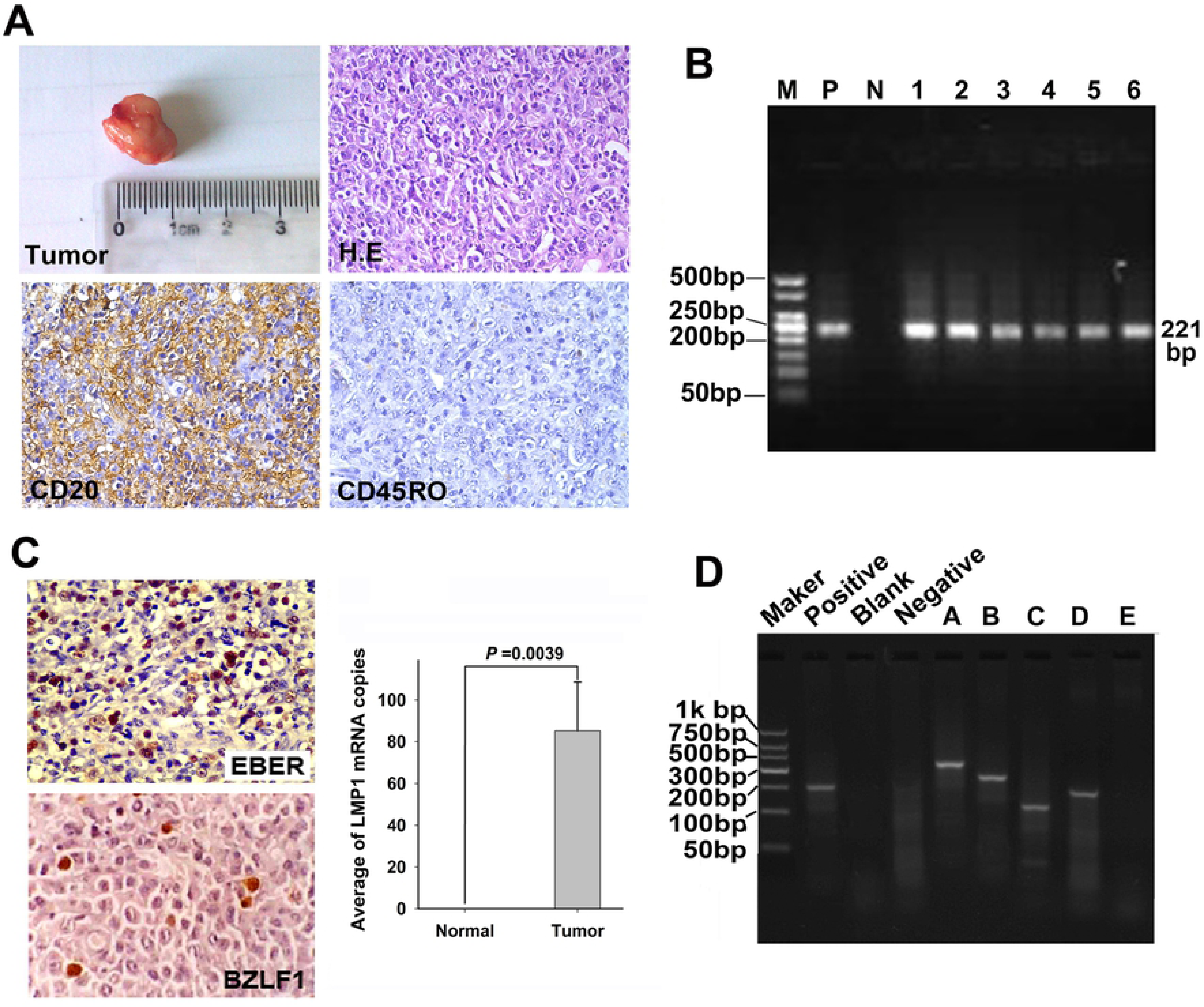
Histopathological and molecular characterization of induced tumors. (A) General morphology of the induced tumor in SCID mice. The tumors were approximately 1.3 × 1.0 × 0.8 cm in size. H&E staining of tumor tissue sections indicated a large tumor cell volume and diffuse large B-cell lymphoma-like morphology. The tumor cells were CD20-positive and CD45RO-negative (×400 magnification). (B) PCR amplification of human-specific *Alu* sequence from the induced tumor. M: DNA marker; P: human PBLs as a positive control; N: mouse organ tissues as a negative control. Lanes 1–6 were induced tumor tissues. A DNA band of 221 bp was clearly visible for all six induced tumors. (C) Detection of EBV expression products in the induced tumors. *In situ* hybridization showed that the tumor cells were positive for EBER with brown nuclei (×400 magnification). A few tumor cells were positive for the expression of the *BZLF1*-encoded protein Zta in the cell nucleus (×400 magnification). Detection by qPCR demonstrated that *LMP1* expression was significantly increased in the induced tumor tissues than in normal PBLs prior to grafting (P< 0.01). (D) Agarose gel electrophoresis of the *IgH* gene fragment amplified from an induced lymphoma by PCR: Lanes A–E represent PCR products amplified from the different primers in the corresponding five tubes. Primers used in each lane are shown in Supplementary Table 2. Analysis of the *IgH* gene rearrangement from the induced tumors showed a single band in lanes A–D, indicating monoclonal growth of tumor cells.

In situ hybridization revealed that the *EBV-encoded small RNA (EBER)* was present in almost all tumor cells and was stained brown in the cell nuclei (Fig 1C). However, tumor-bearing mice organs and tissues adjacent to the induced tumor were negative for *EBER*. In addition, the *BZLF1*-encoded protein *Zta* was positively expressed in the nuclei of the tumor cells. Furthermore, qPCR analysis demonstrated that *LMP1* expression was significantly elevated in the induced tumor tissues than in normal PBLs prior to grafting (p <0.01) (Fig 1C). *IgH* gene rearrangement analysis of the *EBV*-induced lymphomas confirmed that all six induced tumors were derived from the monoclonal proliferation of tumor cells (Fig 1D).

### Analysis of gene expression profiles of the *EBV*-induced lymphomas

Differential gene expression of the *EBV*-induced lymphomas in *SCID mice* and the corresponding normal human lymphocytes were detected with the Agilent human whole genome microarray, and analyzed by SAM, BRB, and LIMMA, respectively. Genes that were identified as differentially expressed were compared across all three methods. A final total of 202 differentially expressed genes were obtained, comprising 44 up-regulated genes and 158 down-regulated genes. Cluster analysis indicated that the 202 differentially expressed genes resulted in good separation of the two sample types (induced lymphoma and normal human lymphocyte from donor). Hence, the genes were considered to be significantly differentially expressed among the two sample types (Fig 2).

**Fig2.**
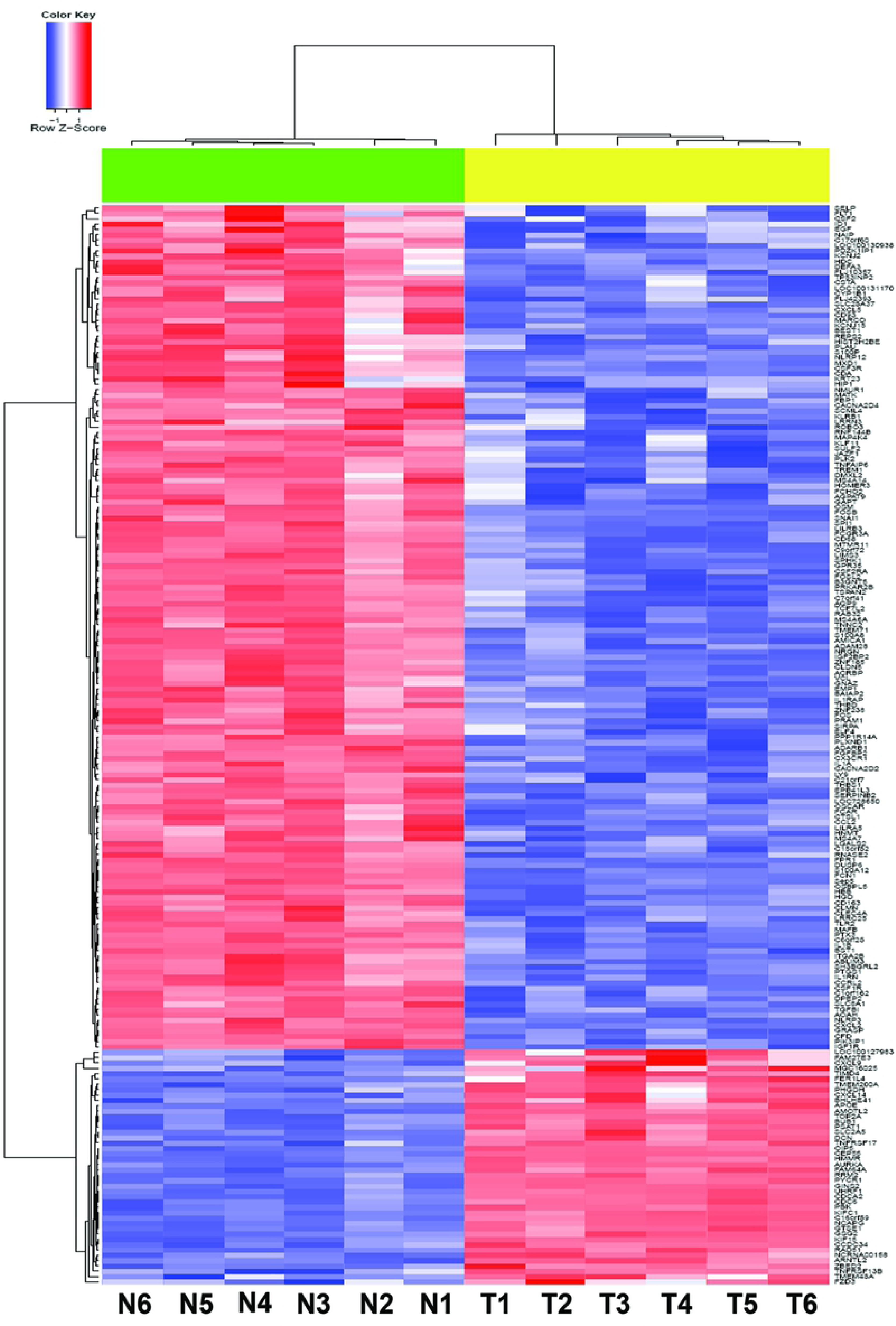
Heat map of differentially expressed genes in EBV-induced lymphomas. Samples were clustered at the top of the tree diagram, with differentially expressed genes clustered on the left side of the diagram. N: Normal human PBLs from the donors, T: EBV-induced lymphoma tissues. Up-regulated genes are shown in red and down-regulated genes are shown in blue. Cluster analysis showed that the 202 differentially expressed genes were mainly clustered into two branches (N and T), demonstrating that these differentially expressed genes can provide good distinction between normal human PBLs and induced lymphomas.

Since these differentially expressed genes interact with other genes to influence relevant cell biological processes, the PPI network was applied to determine the importance of these genes. PPI data that were verified in highly reliable studies were downloaded from the HIPPIE database. By selecting the genes (or proteins) that directly interact with the differentially expressed genes (or proteins), a final total PPI network consisting of 1986 genes (174 differentially expressed genes) and 35280 pairs of interactions were obtained. The importance of a gene in the PPI network can be assessed by various topological properties. In this study, a RS value was calculated for each gene based on the degree, closeness centrality, and clustering coefficient of the gene. The 174 differentially expressed genes were ranked according to RS values. A PPI sub-network was constructed from genes with the top 50 RS values (Fig 3A). Differentially expressed genes with the top ten RS values were *DNA topoisomerase II alpha* (*TOP2A*), *ubiquitin like with PHD and ring finger domains 1* (*UHRF1*), *histone cluster 2 H2B family member E (HIST2H2BE), phosphoglycerate dehydrogenase (PHGDH), vinculin* (*VCL*), *insulin-like growth factor 1 receptor (IGF1R), Fos proto-oncogene (FOS), snail family transcriptional repressor 1 (SNAI1), PDZ Binding Kinase (PBK)*, and *ring finger protein 144B (RNF144B)*. In particular, *TOP2A, UHRF1, PHGDH* and *PBK* were up-regulated, while *HIST2H2BE, VCL, IGF1R, FOS, SNAI1* and *RNF144B* were down-regulated in *EBV*-induced lymphomas. Functions of these genes were mainly enriched in the functions of immune response, inflammation, injury response, defense and chemotaxis, and were all associated with immune or inflammatory responses.

**Fig3.**
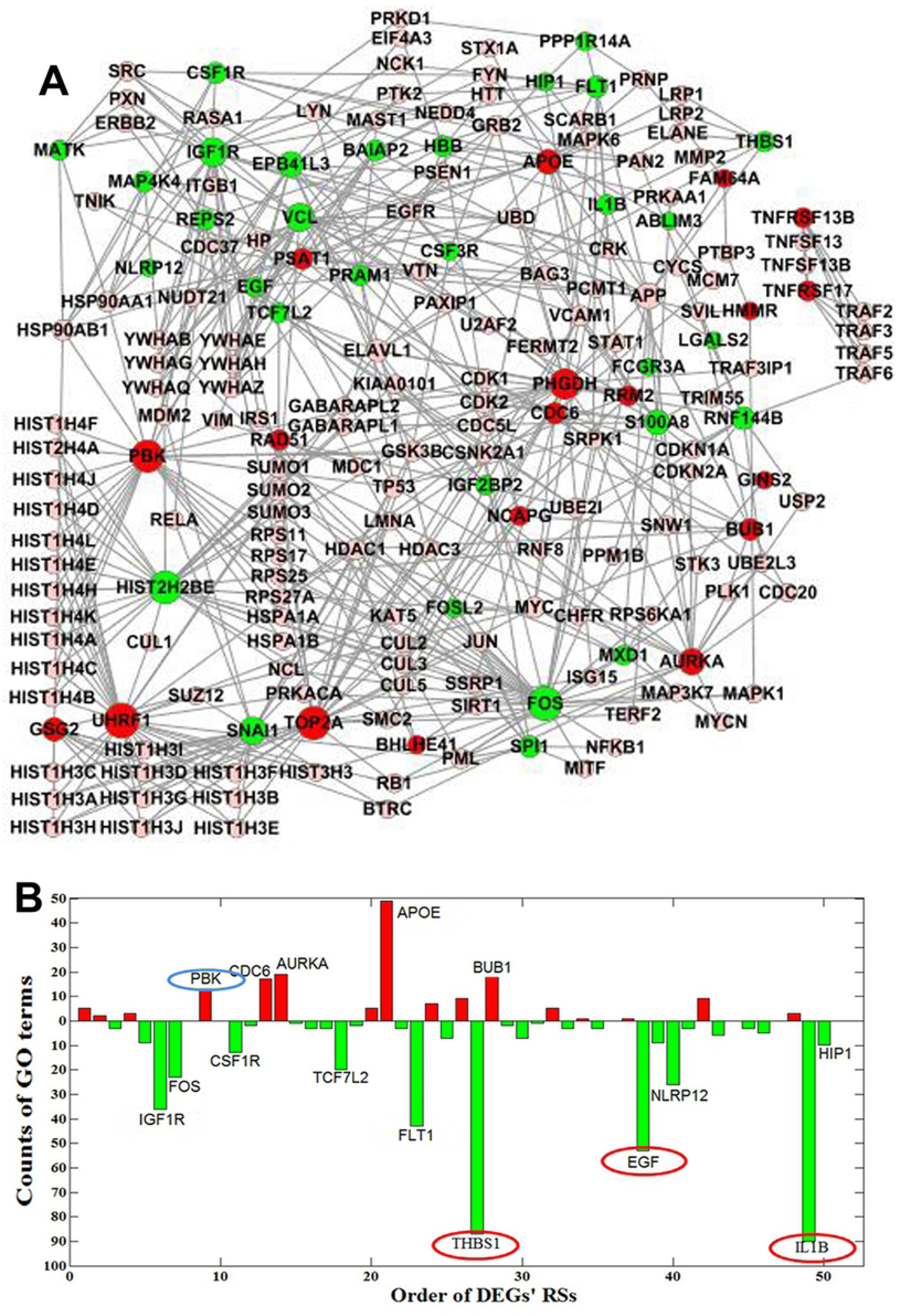
PPI network and comprehensive GO term analysis of the top 50 genes by RS value. (A) PPI network of the top 50 differentially expressed genes by RS value: up-regulated genes are shown in red, down-regulated genes are shown in green, and genes with no significant change in expression are shown in pink. The size of the colored gene dots is related to the number of genes that directly interact with the gene of interest. The higher the number of interacting genes, the larger the dots. (B) Number of GO terms assigned to the top 50 differentially expressed genes by RS value: up-regulated genes are shown in red, down-regulated genes are shown in green. The x-axis indicates the ranking of the differentially expressed gene by RS value, and the y-axis indicates the number of GO terms to which the gene is assigned. Of the top 50 differentially expressed genes by RS value, three with the most GO term annotations are IL-1B, THBS1, and EGF, which are down-regulated in EBV-induced lymphoma. This figure shows that PBK was up-regulated and had the most GO term annotations among the top 10 differentially expressed genes.

Furthermore, gene ontology (GO) enrichment analysis of the differentially expressed genes was performed using the online bioinformatic tool DAVID to calculate the number of GO terms to which the top 50 differentially expressed genes were assigned and to comprehensively assess these genes in combination with the respective RS values (Table 1). GO functional annotations of the top 50 differentially expressed genes by RS value are shown in Fig 3B. The top three genes with the most GO term assignments were *interleukin (IL)-1β, thrombospondin-1 (THBS1)*, and *epidermal growth factor (EGF)* (90, 87, and 53 GO terms, respectively), which were all down-regulated in *EBV*-induced lymphoma. Among the top 10 differentially expressed genes by RS value, the up-regulated gene *PBK* was assigned to the greatest number (13) of GO terms.

### RT-qPCR verification of differentially expressed genes

The mRNA levels of *PBK, IL-1β, EGF, THBS1*, and the reference gene *GAPDH* in the six induced lymphomas and normal PBLs were measured by RT-qPCR. The results showed that the fold change in *PBK* expression was increased in the tumor tissue relative to the control (expression of reference gene was set to 1 after correction), whereas the fold change in the expression levels of *IL-1β, EGF*, and *THBS1* were decreased in the tumor tissue relative to the control. These findings were also consistent with those from the gene expression microarrays (Table 2).

### Functional identification of the candidate key gene *PBK*

Since *PBK* is an up-regulated gene with the most GO terms among the top 10 differentially expressed genes by RS value, the function of *PBK* was further verified using various cell assays.

Firstly, *PBK*-siRNA cell line was first constructed and identified. Upon *PBK*-siRNA-containing lentivirus infection of Daudi cells, cells that were selected by puromycin with stable *PBK* interference were named Daudi/*PBK*-siRNA cells. While *PBK* expression was essentially inhibited in these Daudi/*PBK*-siRNA cells, *PBK* expression was relatively high in Daudi cells as shown by Western blot. The same trends were also observed by RT-qPCR (Fig 4 A, B). The proliferative properties of the Daudi/*PBK*-siRNA cells were then examined. Soft agarose assay analysis of colony formation revealed that the growth of these cells were anchorage-independent and the *in vitro* colony-forming capability of the Daudi/*PBK*-siRNA cells was attenuated, as compared with that of the parental Daudi cells (Fig 4C,D). The viability of the three groups of cells was determined by Cell Titer-Glo. Cell proliferation curves showed that the proliferation of Daudi/*PBK*-siRNA was significantly slowed upon silencing of *PBK* expression (P<0.01) (Fig 4E).

**Fig4.**
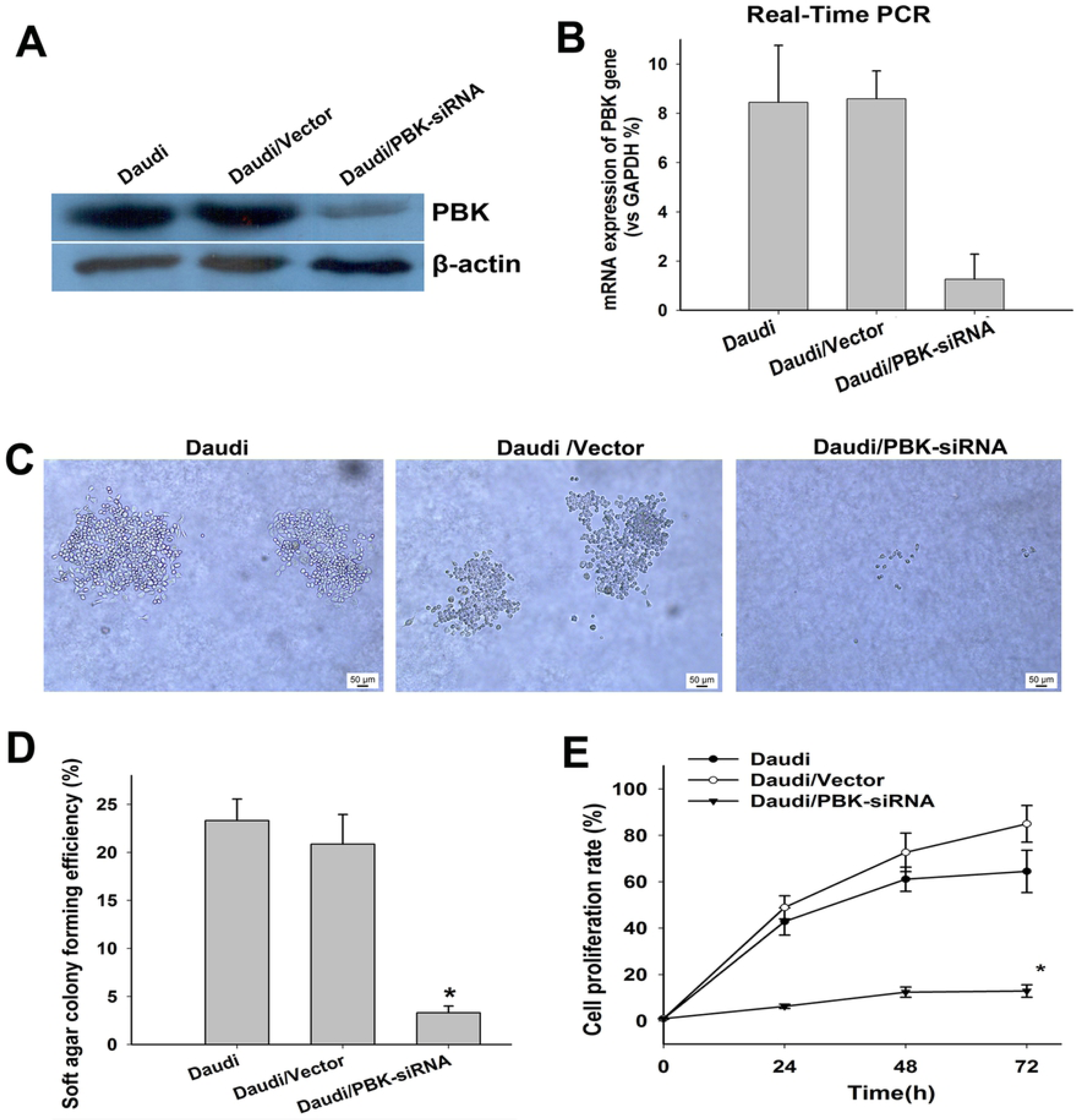
Effect of PBK expression interference on the growth and colony formation of Daudi cells. (A) PBK expression in Daudi cells was knocked down by siRNA and confirmed by Western blot with Daudi/vector as the empty vector control group. (B) PBK mRNA expression levels in Daudi and Daudi/PBK-siRNA cells were determined by RT-qPCR. (C) Daudi cells in each group were evenly inoculated between the double layer agars and incubated for about 2 weeks in serum-free culture medium. Cells were counted under an inverted microscope and the number of colonies and rate of colony formation were recorded based on the >50 cells criteria. (D) Colony formation rate of Daudi cells in the soft agar was calculated and plotted into a column chart for each group. (E) Daudi and Daudi/PBK-siRNA cells were cultured in a 96-well plate, and cell viability detection at 24, 48, and 72h showed that PBK interference inhibited the growth of Daudi cells (\**P*<0.01).

### *PBK* expression in *EBV*-transformed lymphoblasts

Human PBLs were isolated from the peripheral venous blood of seven healthy donors using the lymphocyte isolation solution and were inoculated into a 24-well plate in complete culture medium containing cyclosporine A and *EB virus* solution. After one week of culture, lymphocytes began to increase in size and brightness, and some had grown in clusters (lymphoblast). Then, the cells were divided into additions wells and further cultured. After one month of culture, the number of cells had significantly increased and the cells became overly crowded. These cells were enlarged, translucent, and bright with increased clustering, and were then transferred into culture flasks for further culture. Continued growth of the cells indicated that the *in vitro EBV*-transformed lymphoblast cell line was constructed successfully (Fig 5A).

**Fig5.**
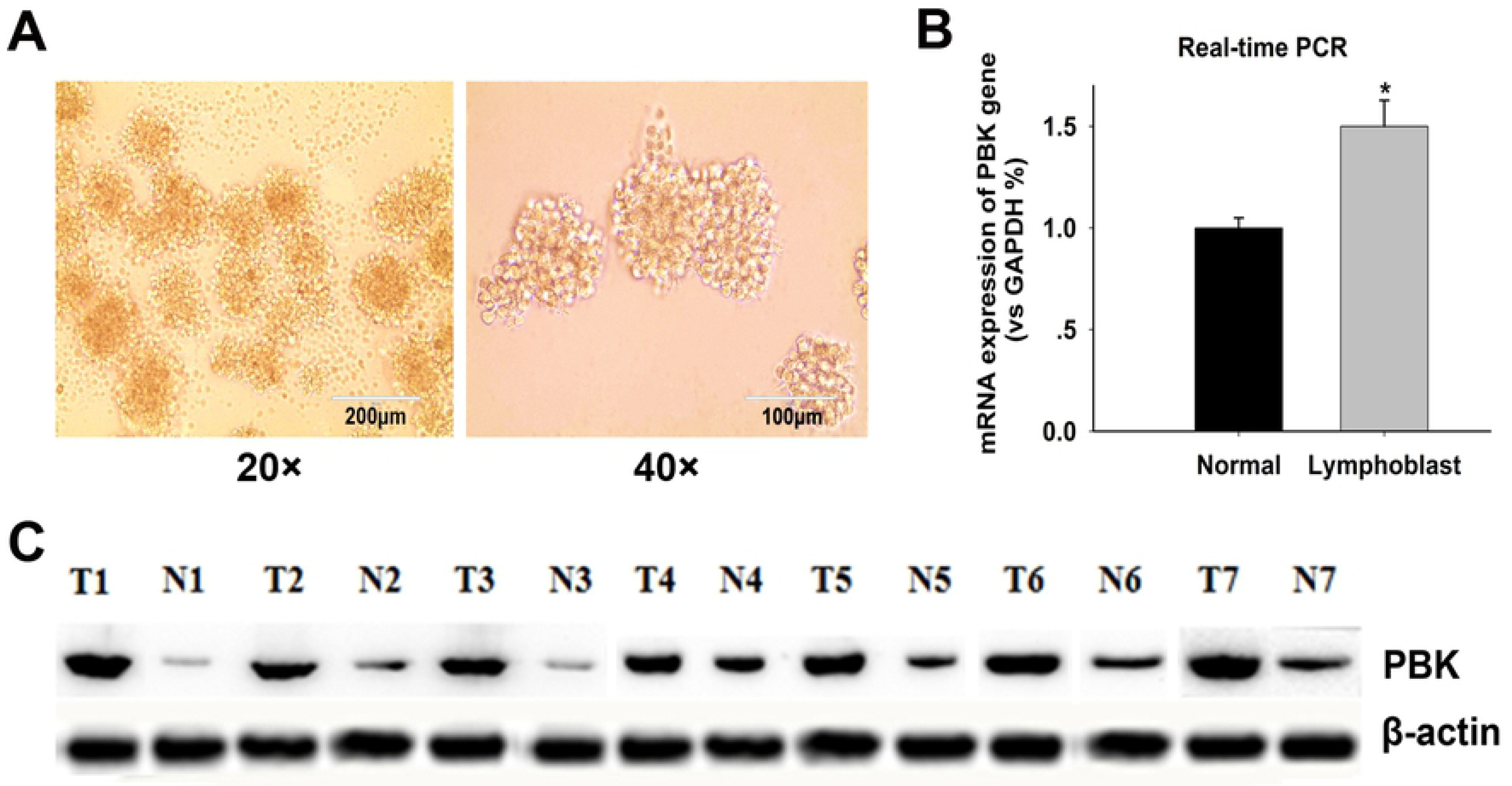
PBK is highly expressed in EBV-transformed lymphoblasts. (A) Microscopic observation showed that EBV-transformed lymphoblasts *in vitro* were relatively enlarged, round, and translucent. The cells were grown in suspension and in clusters, and were capable of continuous passaging. (B) The PBK mRNA levels of EBV-transformed lymphoblasts and normal PBLs were detected by RT-qPCR (\**P*<0.05); (C) PBK protein expression in seven cases of EBV-transformed lymphoblasts (T) and the corresponding normal human PBLs (N) were determined by Western blot.

*PBK* RNA and protein expression of *EBV*-transformed lymphoblasts *in vitro* and the corresponding normal human PBLs were detected by RT-qPCR and Western blot, respectively. The results showed that the *PBK* mRNA and protein levels were elevated in *EBV*-transformed lymphoblasts, as compared to normal PBLs (Fig 5B, C).

### Expression of *PBK* protein in clinical lymphoma specimens

*PBK* protein was positively stained in the nucleus and/or cytoplasm of tumor cells in 49.8% (101/203 cases) malignant lymphoma tissue sections (Fig 6). In particular, *PBK* protein expression was detected in 56.1% (74/132) of B cell lymphomas, 41.5% (22/53) of T cell lymphomas, and 27.8% (5/18) of Hodgkin lymphomas. *PBK* protein expression was identified in 18.2% (8/44 cases) of the reactive lymph node hyperplasia tissues and was mainly distributed in germinal center cells in the follicles of the hyperplastic lymph nodes (Fig 6 D2). Lymphocytes that were adjacent to the follicles were negative for *PBK* protein expression. *PBK* protein expression was significantly higher in lymphomas than in reactive lymph node hyperplasia (p <0.05).

**Fig6.**
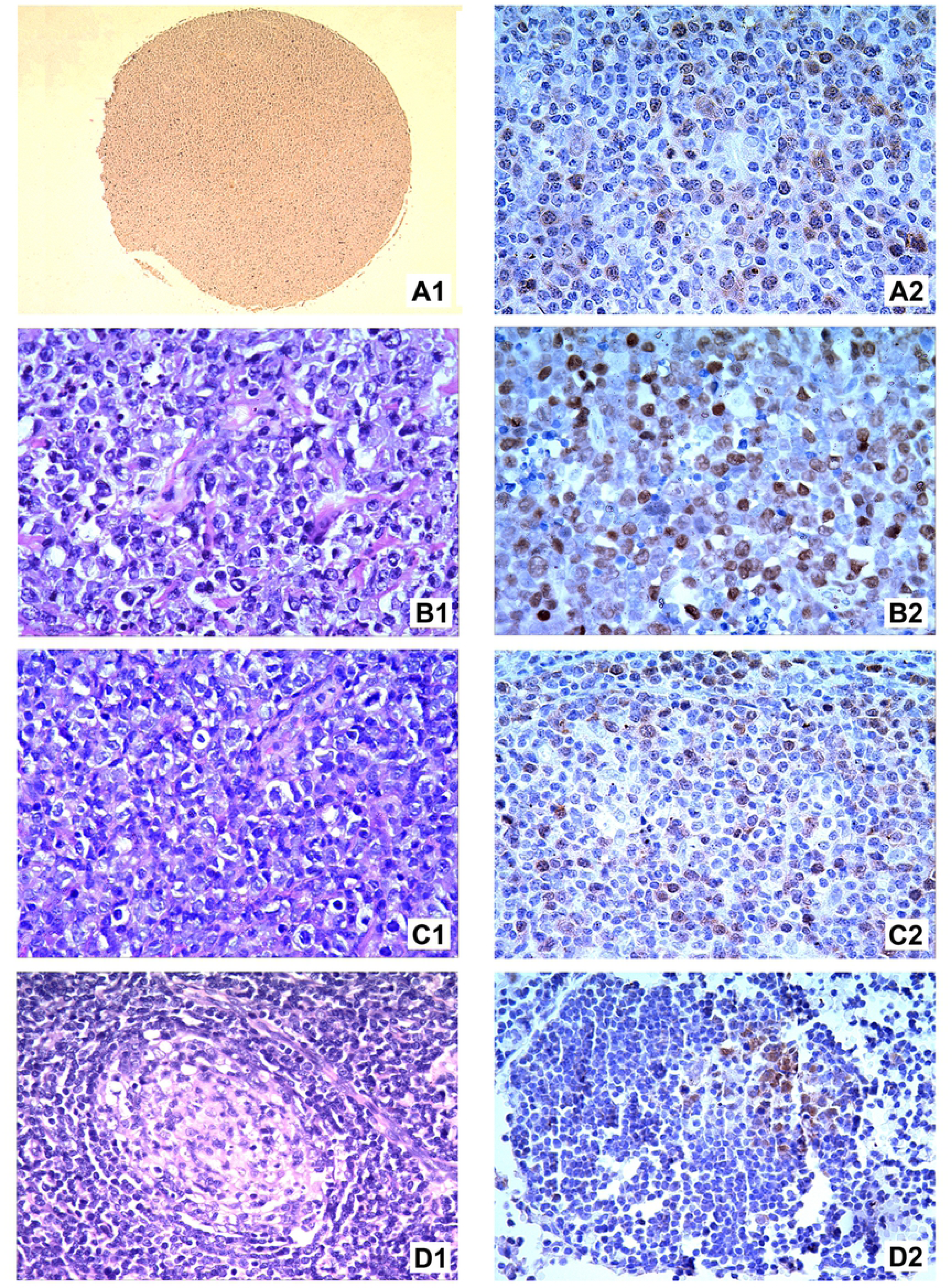
PBK protein expression in clinical lymphomas and reactive lymphnode hyperplasia tissues. A1: Hodgkin lymphoma tissue microarray, SP 4×. A2: Hodgkin lymphoma was positive for PBK, which was distributed throughout the cell nucleus and cytoplasm, SP 40×. B1: B cell lymphoma, H&E 40×; B2: Lymphoma cells were positive for PBK, which was distributed throughout the cell nucleus and cytoplasm, SP 40×. C1: T cell lymphoma, H&E 40×; C2: Lymphoma cells were positive for PBK, SP 40×. D1: Reactive lymphnode hyperplasia, H&E 40×; D2: Germinal center cells in the follicles of the hyperplastic lymph nodes were positive for PBK, SP 40×.

## Discussion

*EBV* is an important human oncogenic virus that is classified as a group 1 carcinogen by the International Agency for Research on Cancer. Since humans are the natural hosts of *EB virus*, there is no ideal experimental animal model to study the relationship between *EBV* infection and tumorigenesis. In order to confirm that *EBV* infection induce the malignant transformation of human lymphocytes and tumor development, normal human PBLs were grafted into *SCID mice* to construct the *hu-PBL/SCID chimeric mice* in our previous study, and showed that *EBV* infection in these chimeric mice could induce the development of human-derived B cell lymphomas [11,12], which was confirmed as a monoclonal proliferative neoplasm by *IgH* gene rearrangement analysis[13]. Based on this previous work, gene expression profiles in lymphoma cells and the corresponding normal donor PBLs were detected in the present study by high-throughput whole genome expression microarray. Differential expressed genes were analyzed using SAM [14], BRB [15] and LIMMA[16], which identified a total of 202 significantly differentially expressed genes, comprising 44 up-regulated genes and 158 down-regulated genes in the *EBV*-induced lymphomas.

A large number of differentially expressed genes are usually generated by gene expression microarray detection. Therefore, the data from a few mainstream PPI databases, such as HIPPIE [17] and BIOGRID [18], were used to determine the PPI network among the differentially expressed genes or between the differentially expressed genes and other relevant genes in order to ultimately elucidate the core positions that the key genes hold within the network. The total PPI network was obtained by mapping the 202 differentially expressed genes identified in this study onto the HIPPIE database. These differentially expressed genes were then ranked by their RS values, which were calculated based on the three topological parameters of each gene in the total PPI network. The top 50 genes by RS values were selected for the construction of a PPI sub-network (Fig 3). The top ten differentially expressed genes based on RS values were *TOP2A, UHRF1, HIST2H2BE, PHGDH, VCL, IGF1R, FOS, SNAI1, PBK* and *RNF144B*. Functional enrichment analysis revealed that these genes were mainly involved in the function of immune response, inflammation, injury response, defense, and chemotaxis, which are all associated with the immune or inflammatory responses.

GO term analysis of the differentially expressed genes using the online bioinformatic software DAVID showed that the top three genes with the highest number of GO terms among the top 50 genes ranked by RS were *IL-1β, THBS1*, and *EGF* (90, 87, and 53 GO terms, respectively). These genes were all down-regulated in the lymphomas. On the other hand, the up-regulated gene *PBK* has the highest number (13) of GO terms among the top 10 genes ranked by RS.

A study of the differentially expressed genes in *EBV*-transformed lymphoblastoid cell lines and the corresponding normal B cells by Caliskan identified a total of 2,217 highly expressed differential genes in lymphoblastoid cell lines [19]. GO analysis revealed that these genes were involved in various biological functions, including cytokine activity, signal transduction and the immune response. This finding was similar to that of the GO analysis in our study. Further analysis by expression quantitative trait loci identified 160 *EBV* infection-associated genes, which included 35 involved in the regulation of anti-apoptotic signals in the NF-κB pathway, and hence have added a new bioinformatic evidence to the existing consensus [20,21]. Zhang et al. detected the gene expression profiles of seven diffuse large B cell lymphomas and seven normal lymph node tissues using whole genome expression microarray, and obtained 945 differentially expressed genes comprised of 272 up-regulated genes and 673 down-regulated genes [22]. Enrichment analysis using the Kyoto Encyclopedia of Genes and Genomes (KEGG) showed that these genes were significantly enriched in two pathways, namely the interactions between immune functions and signaling molecules. In our present study, the differentially gene expression profiles of *EBV*-induced lymphomas and parental lymphocytes in *human PBL/SCID chimeric mice* were analyzed, which identified key differentiated genes related to cell growth and differentiation, and the immune response. Then, the up-regulated gene *PBK* with the most GO terms among the top 10 genes ranked by RS values in the PPI network was selected for further study of its effect on cell growth and proliferation.

*PBK*, also known as *TOPK*, is a serine/threonine kinase in the dual specificity mitogen-activated protein kinase family. *PBK* can induce the activation of lymphoid cells by destabilizing tumor suppressor p53 as it becomes phosphorylated, which in turn facilitates the transition through the G2/M checkpoint [23]. Previous studies have shown that *PBK* expression was significantly higher in diffuse large B cell lymphomas than in normal lymphocytes [24], and up-regulated in some malignancies [25,26,27], indicating that *PBK* may play a critical role in tumor development. In this study, both Western blot and qPCR analyses demonstrated that *PBK* expression was higher in *EBV*-transformed lymphoblasts *in vitro* and clinical lymphoma specimens than in normal human lymphocytes, which was also consistent with the finding by the gene expression microarray, suggesting that *EBV* may up-regulate the expression of *PBK* in B cells via activation of some signaling pathways. A study by Chen et al. found that the transcription factor *E2F-1* can directly promote the transcription of *PBK* [28], and immediate-early protein *BZLF1* of *EB virus* can up-regulate the expression of *E2F-1* [29,30,31]. Interestingly, *BZLF1* expression was also detected in *EBV*-induced lymphomas in our study, thus we speculated that the *BZLF1*→*E2F-1*→*PBK* positive regulatory pathway may partially explain the high expression of *PBK* in *EBV*-induced B cell lymphoma. To verify the function of *PBK, PBK* expression was silenced in *EBV*-positive Daudi cells, which showed that cell growth, proliferation, and colony formation *in vitro* were significantly attenuated upon *PBK* silencing, which further indicated that *PBK* may have an important role in the *EBV*-facilitated malignant transformation of human B cells.

Of the top 50 genes ranked by RS values, the top three with the most GO terms were *IL-1β, THBS1*, and *EGF. IL-1β* is a cytokine associated with immune regulation and tumor cell immunoevasion that is involved in various immune responses, and hence an important mediator of the immune system. *IL-1β* is synthesized as a precursor by monocytes or macrophages, and is modified and released by enzymatic cleavage during cell damage to induce the apoptosis of damaged cells within the local tissue [32]. Studies have found that *IL-1β* can inhibit the growth and metastasis of tumor cells via the host response [33], and the secreted *IL-1β* receptor antagonists have similar biological effects [34], indicating that *IL-1β* may be a tumor-suppressor in the tumor microenvironment. Our microarray analysis revealed that *IL-1β* was significantly down-regulated in *EBV*-induced lymphomas, as compared to the control, suggesting that *EBV* infection may affect the expression and activity of this cytokine in the tumor microenvironment through several mechanisms, and thereby permit immunoevasion of *EBV*-infected lymphocytes. The *THBS1* protein is synthesized by endothelial and immune cells, and can affect cell phenotype and extracellular matrix structure via glycoproteins, and participate tumor angiogenesis and tissue remodeling [35]. *THBS1* can activate transforming growth factor-β to inhibit the migration, growth, and survival of vascular endothelial cells in the tumor microenvironment, and thereby inhibit tumor progression [36]. We found that *THBS1* expression was down-regulated in *EBV*-induced lymphomas, suggesting that the signaling network in *EBV*-induced lymphomas tends to promote the growth of vascular endothelial cells, which facilitate the development and progression of tumors. In many cancers, *EGF* can activate receptor tyrosine kinases by binding to c-erb-B receptor families and regulate the proliferation, differentiation, survival, angiogenesis, and migration of cancer cells [37,38]. However, further studies will be needed to elucidate the role and mechanism of *IL-1β, THBS1*, and *EGF* down-regulation in *EBV*-induced lymphoma.

In this study, we obtained the gene expression profile of *EBV*-induced lymphomas in an animal model. By constructing the gene-function relationship network, we preliminarily found that *TOP2A, UHRF1, HIST2H2BE, PHGDH, VCL, IGF1R, FOS, SNAI1, PBK* and *RNF144B* may be the key genes in *EBV*-induced lymphoma. In particular, *PBK* is an important key node gene that may become a new molecular target for the prevention and treatment of human *EBV*-associated lymphomas.

## Materials and Methods

### Donors and animals

Peripheral venous blood samples (300∼400 mL) collected from healthy donors were provided by the Hengyang Central Blood Station. Male and female *SCID mice* (NOD. CB17-Prkdc scid /NcrCrlVr) at 3∼4 weeks of age, weighing 18.00 ± 0.53 g were purchased from Beijing HFK Bioscience Co., Ltd. (Beijing, China; license no. SCXK2009-0004; Certificate No. Jing 0212751). All experiments were approved by the Research Ethics Committee of Medicine, University of South China.

### Reagents and antibodies

Agilent whole genome microarrays (Agilent Technologies, Santa Clara, CA, USA) were used for gene expression profile detection. The RNeasy Mini Kit was purchased from Qiagen (Hilden, Germany; catalogue no. 74104), the Quick Amp Labeling Kit (single label) was purchased from Agilent Technologies (version 5.7, catalogue no. 5190-0442), the Gene Expression Hybridization Kit was purchased from Agilent Technologies (catalogue no. 5188-5242), and the Q-PCR kit was purchased from Tiangen Biotech Co., Ltd., (Beijing, China; catalogue no. FP207).

Monoclonal antibodies against human lymphocytes (LCA, CD45), B cells (L26/CD20, CD79a) and T cells (CD45RO, CD3) were purchased from Maxin Co., San Francisco, LA. Antibody against *PDZ Binding Kinase* (*PBK*)/*T-LAK cell-originated protein kinase (TOPK)* was purchased from Cell Signaling Technology, Inc. (Beverly, MA, USA; #4942).

### Cell culture and tissue sample

Daudi cells were purchased from the China Center for Type Culture Collection (Wuhan, China) and 293FT cells were stored in our laboratory. Bovine serum albumin and the Cell Titer-Glo Luminescent Cell Viability Assay Kit were purchased from Promega Corporation (Madison, WI, USA).

Daudi and 293FT cells were cultured in complete Roswell Park Memorial Institute (RPMI)-1640 culture medium containing 10% fetal bovine serum (FBS). *EBV*-transformed lymphoblasts were also cultured in complete RPMI-1640 culture medium.

A total of 113 malignant lymphomas and 44 reactive lymph node hyperplasia paraffin-embedded tissue specimens were collected from Pathology Department of The First Affiliated Hospital and The Second Affiliated Hospital, University of South China. Lymphoma tissue microarrays were purchased from Auragene Bioscience Corporation, Inc. (Changsha, China; product no. TC0009). Each microarray includes 90 lymphoma samples, and each lymphoma sample has been clinicopathologically diagnosed and comes with a complete set of pathologic examination and immunohistochemical staining data.

### Plasmids

Lentiviral vector system includes the pLEX-MCS plasmid and the PMD2G and psPAX2 packaging plasmids (Thermo Fisher Scientific, Waltham, MA, USA).

### Construction of *hu-PBL/ SCID chimeric mice* and *EBV* infection

Peripheral blood lymphocytes (PBLs) were isolated from the fresh blood samples of six healthy donors and intraperitoneally grafted into *SCID mice* at 8∼10 × 10^7^PBLs/mouse. PBLs from each donor were grafted into 3∼4 *SCID mice*.

*EBV* was isolated from the supernatant of B_95-8_ cells cultured *in vitro* under starvation conditions. *EBV* infection was not required for *SCID mice* that were grafted with PBLs from donors who were positive for *EBV* viral capsid antigen (VCA)-IgG and IgA. On the other hand, 0.4 mL of *EBV* suspension was intraperitoneally injected into *SCID mice* that were grafted with PBLs from donors who were VCA-IgA negative on day 3 post-grafting. Details on the *EBV* infection status of the blood donors are shown in Supplementary Table 1.

Food intake and activity of the *SCID mice* were regularly observed, and dying mice were immediately euthanized. All surviving mice were euthanized on day 135 post-PBL grafting. Detailed dissection was performed on all animals. Portions of the fresh tumor tissues were harvested and stored in liquid nitrogen, while the remaining tumor tissues and other major organs were fixed in 10% neutral formalin solution.

### Examination of *EBV*-induced tumors

The tumor tissue sections in *hu-PBL/SCID mice* were stained with hematoxylin and eosin (H&E) for microscopic examination. Expression of several antigens, including human leukocyte common antigen (LCA), B cell markers (CD20 and CD79a), T cell markers (CD3 and CD45RO), and *EBV*-encoded transcriptional activator *Zta* (*BZLF1*), were determined by immunohistochemical (IHC) stain.

The human Alu sequence (221 bp) was amplified by PCR using the primer 5’-CAC CTG TAA TCC CAG CAG TTT-3’, 5’-CGC GATCTC GGC TCA CTG CA-3’.

*EBV-encoded small RNA (EBER)* in tumor tissues was measured by in situ hybridization using the hybridization detection kit (Triplex International Biosciences Co., Ltd., Fuzhou, China; catalogue no. S30172) as per the manufacturer’s instructions. The presence of brownish-yellow granules in the cell nuclei was considered to be positive for *EBV* infection, while the absence was considered to be negative.

Differences in *latent membrane protein 1 (LMP1)* mRNA expression of *EB virus* between the induced tumor tissues and normal lymphocytes was compared using real-time quantitative PCR (qPCR) with the primer 5’-CTG CTC ATC GCT CTC TGG AA-3’. Each sample was run in triplicate using *glyceraldehyde 3-phosphate dehydrogenase* (*GAPDH*) as the reference gene.

For determination of clonality of *EBV*-induced tumors, genomic DNA was extracted from the tumor tissue sections as per the instructions of the DNA FFPE Tissue Kit (Qiagen, catalogue no. 56404). Clinically and pathologically diagnosed lymphoma tissue specimens were used as the monoclonal (positive) control, and a reactive lymph node hyperplasia specimen was used as the polyclonal (negative) control. The human *immunoglobulin heavy chain (IgH)* gene was amplified by PCR using the BIOMED-2 standardized primer system [39]. The PCR reaction was conducted in five tubes labelled A, B, C, D, and E. The qPCR reactions were performed with the primers shown in Supplementary Table 2 under the following conditions: 95°Cfor 7 min, followed by 40 cycles at 95°Cfor 45 s, 60°Cfor 45 s, and 72°Cfor 90 s, followed a final extension cycle at 72°Cfor 10 min. The PCR amplicons (8 μL) were then verified by 2.5% agarose gel electrophoresis and visualized under a ultra-violet transilluminator using the gel documentation imaging system (Vilber LourmatSté, Collégien, France).

### Gene expression microarray detection and analysis

Total RNA was extracted from the six *EBV*-induced lymphomas (tumor group) and normal human lymphocytes (normal group) using TRIZOL reagent (Ambion/Life Sciences, Grand Island, NY, USA), according to the manufacturer’s instructions, and labelled as per the instructions of the Agilent Quick Amp Labeling Kit (Agilent Technologies). First-strand cDNA was synthesized from purified total RNA (template) using the T7 promoter primer and Moloney murine leukemia virus reverse transcriptase, followed by the synthesis of the second-strand cDNA. Both cDNA strands were then synthesized into cRNA using transcription master mix and labelled with cyanine-3-cytidine-5’-triphosphate.

Gene expression profiles of the normal and tumor groups were detected using Agilent whole genome microarray containing 41,000 known human gene transcripts. The microarrays were scanned with the Agilent Microarray Scanner and the raw data were recorded and normalized using Agilent Feature Extraction Software to calculate the signal fold change between the tumor and normal groups. A ≥2-fold change in gene expression, as determined by the t-test, was considered as up-regulated, and a ≤0.5-fold change in gene expression was considered as down-regulated (all differentially expressed genes were p < 0.05).

Differential gene expression was determined using three methods, namely significance analysis of microarrays (SAM) [14], Biometric Research Program (BRB) [15], and Linear Models for Microarray and RNA-Seq analysis (LIMMA) [16], to ensure that the genes were significantly differentially expressed (differential expression indicated by all three methods). The threshold value for all three methods was set at 2-fold change and a false discovery rate of <0.001. Functional enrichment analysis of the differentially expressed genes was conducted using the online bioinformatics tool DAVID (https://david.ncifcrf.gov/) [40].

### Protein-protein interaction (PPI) analysis was subsequently performed

First, the PPI data, as verified in highly reliable studies, were downloaded from the Human Integrated Protein-protein Interaction rEference (HIPPIE) database (http://cbdm.mdc-berlin.de/tools/hippie) [17]and the proteins in the PPI network were then converted into corresponding gene symbols listed in the National Center for Biotechnology Information database (https://www.ncbi.nlm.nih.gov), along with the probes in the microarrays. After acquiring the PPI network, the topological properties of the genes within the network were calculated, including the degree, closeness centrality, and clustering coefficient [41,42,43]. Then, the genes were sorted and scored (mean value) based on these three topological properties using the formula RS_i_=(Rank(k_i_) +Rank(C_i_)) +Rank(CC_i_))/3. Finally, the importance of the genes within the network were assessed based on rank score (RS) values [44]. The topological properties of the network were computed and graphed using the open source software Cytoscape (http://www.cytoscape.org/) [45].

### RT-qPCR detection of differentially expressed genes

Total RNA was extracted from *EBV*-induced tumor cells (tumor group) and normal human lymphocytes (normal group) and reverse transcribed into cDNA. Then, differentially expressed genes were amplified by real-time PCR using the primers *GAPDH*: F:5’-GGGAAACTGTGGCGTGAT-3’,R:5’-GAGTGGGTGTCGCTGTTGAN -3’; *PBK*: F:5’-TTTCCTTCCAGGCGGTGAG-3’, R:5’-ACGGAGAGGCCGGGATA TT-3’; IL1B: F:5’-CGAATCTCCGACCACCACTACA-3’, R:5’-AGGGAACCAGCA TCTTCCTCAG-3’; *EGF*: F:5’-AAAACGCCGAAGACTTACCC-3’, R:5’-AACCTTC ACGACACGAACACC-3’; *THBS1*: F:5’-TGGAAAGATTTCACCGCCTAC-3’, R:5’-CCTGGGGGTTTTCTCAAGC-3’ under the following conditions: 95°Cfor 10 min, followed by 40 cycles at 94°Cfor 10 s, 61°Cfor 20 s, and 72°Cfor 25 s. *GAPDH* was used as the reference gene. Melting curve analysis was performed using the default settings of the instrument. Relative gene expression was calculated according to the 2^- ΔCCT^ method.

### Functional experiment of *PBK* gene

Small interfering RNA (siRNA) against *PBK* was prepared. One negative control and three *PBK*-siRNA-specific antisense oligonucleotide fragments were synthesized by Guangzhou RiboBio Co., Ltd. (Guangzhou, China). The primer for siRNA1, siRNA2, siRNA3 and the siRNA negative control were 5’-AACCGAACUUACCAAGCAU-3’; 5’-ACCCUAAGAACAUGGCCUA-3’; siRNA3: 5’-AUGGCAACCCUAAG AACAU-3’ and 5’-AAATAAGAATGGCCCCAT-3’, respectively.

### Packaging and transfection of *PBK*-siRNA lentiviral particles

Viral packaging was completed by Guangzhou RiboBio Co., Ltd. A total of 5 mL of concentrated *PBK*-siRNA viral particles and 5 mL of empty viral particles were obtained. Infected cells were cultured and selected in Dulbecco’s modified Eagle’s medium (DMEM) containing 10% FBS and 2.0 μg/mL of puromycin. The mRNA and protein levels of the *PBK* gene in Daudi cells were determined by qPCR and immunoblot analysis, respectively, in order to measure the rate of interference.

### Western blot detection

Total protein was extracted from Daudi cells in the log phase using RAPI assay protein lysis buffer, and the concentration was determined with the bicinchoninic acid assay. The proteins (70 μg) were separated by sodium dodecyl sulfate polyacrylamide gel electrophoresis on 10% gels and then transferred onto a polyvinylidene difluoride membrane, which was then blocked and incubated with the primary antibody, followed by a horseradish peroxidase-conjugated secondary antibody. The membrane was visualized after incubation with enhanced chemilumescentsubstrate.

### Detection of cell proliferation with the Cell Titer-Glo luminescent cell viability assay

Daudi cells in the log phase were seeded at 1 × 10^4^ cells/well in triplicate wells of a 96-well plate in a total volume of 180 μL. After culturing the cells until the death phase, culture media was removed from the wells and the cells were washed three times with phosphate-buffered saline, followed by the addition of 100 μL of detection reagent into each well. After gently shaking the plate for 2 min, the cells were incubated for 10 min in the dark, and proliferation was then measured on the instrument under the detection parameters of scintillation counter mode, distance above the well of 1 mm, and count for 10 s. The experiment was repeated three times.

### Soft agar assay for colony formation

Daudi cells in the log phase were prepared into a single-cell suspension and adjusted to a concentration of 1 × 10^6^cells/L in DMEM culture medium containing 20% FBS. Two-layered low melting point agarose was prepared as per the methods in a previous study [46]. The cells were mixed with the top agarose layer, incubated at 37°Cfor 10∼14 days under an atmosphere of 5% CO_2_, and then observed under an inverted microscope to calculate the number of clones and rate of clone formation based on the presence of >50 clones in a colony.

### Detection of *PBK* expression in *EBV*-transformed lymphocytes

A total of 1.5 mL of *EB virus* solution and 2 mL of complete culture medium were added to centrifuge tubes containing the lymphocyte pellets from the blood donors. The cell pellets were resuspended thoroughly. Then, each mixture was equally divided into two wells in a 24-well plate and cultured at 37°Cunder an atmosphere of 5% CO_2_. After 7 days of culture, the cells were observed under a microscope, which showed significantly enlarged and brighter lymphocytes, and the formation of cell clusters (lymphoblasts). At this time, 1/3 of the culture medium was replaced with fresh medium and 1/2 of the culture medium was replenished every 3∼4 days thereafter. On day 15, the number of lymphoblasts was significantly increased and the cells were divided into four wells. As the cells proliferated and became crowded, they were further divided into eight wells or transferred to a 50-mL culture flask for further culture. At this time, the cells should appear large, translucent, and bright. After 6∼8 weeks of culture, the cells were collected and frozen for 2 weeks and then thawed again for culture. If the cells were able to continue growing, then an *in vitro EBV*-transformed lymphoblastoid cell line was considered to be established successfully.

*PBK* expression of the *EBV*-transformed lymphoblasts was measured by RT-qPCR. Forward and reverse primers for *PBK* were same as above.

### Detection of *PBK* protein expression in malignant lymphomas

Immunohistochemical staining was performed on a total of 203 cases of clinic tissue specimens, including 90 lymphoma samples from the microarray and 113 resected lymphoma samples, as per the instructions of the SP kit using the *PBK* antibody (Cell Signaling Technology, Inc.;#4942).

### Statistical analysis

All statistical analyses were performed using IBM SPSS Statistics for Windows, version 19.0 (IBM Corp., Armonk, NY, USA). Quantitative data are expressed as the mean ± standard deviation (X ±SD). A probability (p) value of<0.05 was considered as statistically significant.

## Acknowledgments

This work was supported by the National Natural Science Foundation of China (No. 81372134,81641012 and 81500169), Hunan Province Key Laboratory of Tumor Cellular & Molecular Pathology (2016TP1015), and Hunan Province Cooperative Innovation Center for Molecular Target New Drug Study (2014-405).

## Author Contributions

**Conceptualization:** Yimou Wu, Runliang Gan

**Data curation:** Chengkun Wang, Runliang Gan.

**Formal analysis:** Yang Zhang, Xiaoning Gan, Lu Xie, Chengkun Wang.

**Funding acquisition:** Yang Zhang, Runliang Gan, Yunlian Tang.

**Investigation:** Yang Zhang, Rongfang He.

**Methodology:** Chengkun Wang, Qiu Peng, MeiyingTan, Runliang Gan.

**Project administration:** Yang Zhang, Yimou Wu.

**Supervision:** Runliang Gan.

**Writing – original draft:** Yang Zhang, Chengkun Wang.

**Writing – review & editing:** Yimou Wu, Runliang Gan.

## Supporting Information Legends

